# Spatio-temporal analysis of remote sensing images provides early warning signals of forest mortality

**DOI:** 10.1101/2021.09.18.460897

**Authors:** Sara Alibakhshi

## Abstract

Ecosystems are under unprecedented pressures, reflected in rapid changes in the regime of disturbances that may cause negative impacts on them. This highlights the importance of characterizing the state of an ecosystem and its response to disturbances, which is known as a notoriously difficult task. The state-of-the-art knowledge has been tested rarely in real ecosystems for a number of reasons such as mismatches between the time scale of ecosystem processes and data accessibility as well as weaknesses in the performance of available methods. This study aims to use remotely sensed spatio-temporal data to identify early warning signals of forest mortality using satellite images. For this purpose, I propose a new approach that measures local spatial autocorrelation (using local Moran’s I and local Geary’s c method) at each time, which proved to produce robust results in multiple different study sites examined in this article. This new approach successfully generates early warning signals from time series of local spatial autocorrelation values in unhealthy study sites 2 years prior to forest mortality occurrence. Furthermore, I develop a new R package, called “stew”, that enables users to explore spatio-temporal analysis of ecosystem state changes. This work corroborates the suggestion that spatio-temporal indicators have the potential to diagnose early warning signals to identify upcoming climate-induced forest mortality, up to two years before its occurrence.

## 1. Introduction

Ecosystems are under unprecedented pressures, reflected in rapid changes in the environment and regime of disturbances (Veraart et al., 2012). Such pressures threaten the resilience of an ecosystem, increasing the risk of irreversible critical transitions and pushing the state of an ecosystem to an alternative state (Scheffer et al., 2001). Critical transitions have been already observed in a variety of ecosystems, e.g., a transition of a wetland to rangeland (Alibakhshi et al., 2017), or transition of a forest to a savanna or a treeless state (Hirota et al., 2011). Under human pressures, the likelihood of such critical transitions is expected to increase in ecosystems (Veraart et al., 2012) and it suggests the necessity to improve our understanding of ecosystem feedback upon disturbances.

Critical transitions may occur when the resilience of an ecosystem following a disturbance (the ecosystem capacity to recover from disturbances) reduces, leading to a slow recovery of ecosystem function. Under such circumstances, the ecosystem may approach a critical threshold (tipping point) where even a small perturbation may cause a large response, called a critical transition. This highlights the need for developing and enhancing the approaches for identifying the dynamics of ecosystems, however, numerical approaches for its quantification are developing slowly (Angeler and Allen, 2016).

Quantifying resilience and predicting whether, where, and when an ecosystem may experience critical transition, are a notoriously difficult task (Sato and Lindenmayer, 2018; Scheffer et al., 2001). This is because most of the ecosystems are highly dynamic and complex with non-linearity and peculiarities of processes interacting with multiple driving factors varying in space and time. To tackle the difficulties, recent studies suggested a set of methods, called early warning signals (or so-called leading indicators) of critical transitions. These methods use statistical metrics such as lag-1 autocorrelation, standard deviation or mathematical approaches on a time series of data (Scheffer et al., 2009). These indicators rely on the “critical slowing down” theory which implies that the resilience of an ecosystem upon disturbances becomes slower when the ecosystem approaches a tipping point (Scheffer et al., 2009). In this theory, the loss of resilience may be inferred as early warning signals of a possible upcoming critical transition or abrupt change in the state of an ecosystem (Dakos et al., 2015). The implication of these methods in this study is, for example, to test whether the reduced resilience can be captured timely and how they are linked to forest mortalities.

Using early warning signals (or so-called leading indicators) of critical transitions, we can explore the state of an ecosystem by quantifying key ecosystem variables (hereafter, ecosystem state variable) (Dakos et al., 2012a). Ecosystem state variables have the property to represent the state of an ecosystem directly or indirectly. For example, the normalized difference vegetation index (NDVI) might be a proper leading indicator that has a potential to represent the state of a forest ecosystem (Liu et al., 2019; Rogers et al., 2018). Observing, for example, an increase in the trend of the lag-1 temporal autocorrelation of an ecosystem state variable indicates a decline in the resilience of the ecosystem that can be considered as an early warning signals of possible upcoming critical transitions or abrupt changes (Dakos et al., 2012a, 2012b). These early warning signals generated from the temporal analysis are, hereafter, called temporal early warnings (TEW). Some studies also suggested a set of methods which explores the variation of an ecosystem state variable along a spatial gradient instead of time. These methods are applied to spatial data in an ecosystem where alternative states can be measured along a spatial gradient (e.g., transition of the forest to savanna). For instance, increasing a spatial autocorrelation along the a spatial gradient can be considered as an early warning signal of critical transitions (Kéfi et al., 2014).

Although successful applications of critical transitions approaches on space and time have been mostly reported in fully controlled experimental studies or using simulation data (e.g., Carpenter et al., 2011; Carpenter and Brock, 2006), their applications in real ecosystems have been limited. Moreover, they are still subjected to false alarms (either false negative or false positive alarms of upcoming state change in an ecosystem) (Dakos et al., 2015; Nijp et al., 2019). For instance, the methods generating TEW are sensitive to pre-processing steps and their performance may be affected by the size of the data (Dakos et al., 2012a). The spatial indicators are also subjected to the risk of false alarms due to the assumption of homogeneity in a spatial gradient that is usually invalid in ecosystems (Nijp et al., 2019; Radford and Bennett, 2004). In addition, one of the greatest obstacles to the success of these methods was a lack of appropriate data that can timely present the state of an ecosystem (Dakos et al., 2015; Scheffer et al., 2009). In recent years, vast investments have been increasingly dedicated to produce remotely sensed data measuring states and fluxes of various environmental properties in space and time (Reichstein et al., 2019).

Satellite images provide a great opportunity that may enhance our ability to measure an ecosystem’s resilience (Dakos et al., 2015). For instance, the use of such spatio-temporal data allows the indicators to generate TEW on each spatial location (pixel) separately, providing multiple replications of temporal observations across the ecosystem and generates a spatial pattern of TEW (hereafter, SPT). Such a spatial pattern, resulting from multiple replications of temporal observations over different pixels, may provide more reliable early warning signals and decrease the risk of generating false alarms compared with using a single temporal data that represents the entire ecosystem.

This study aims to explore the potentials of spatio-temporal optical satellite images to understand the dynamics of forests and infer whether and where the early signals of reduced resilience and forest mortality can be identified. I hypothesized that multiple replications of temporal data obtained over spatial locations (from spatio-temporal images) can improve capturing early warning signals in forest ecosystems. In addition, I suggest a new approach that uses local spatial autocorrelation statistics (i.e., local Moran’s *I* and local Geary’s *c*), applied to an ecosystem state variable at each time pixelwise, and tested whether the variation of local spatial structure at each pixel over time (hereafter, STM and STG for spatio-temporal variation of local Moran’s *I* and local Geray’s *c*, respectively) can provide early warning signals of forest decline. I expected that the STM and STG methods may improve our ability to identify early warning signals of forest mortality, since these methods are less sensitive to pre-processing steps and independent of the assumption of landscape homogeneity.

To test my hypothesis, I selected three case studies including two unhealthy forests with a known history of reduced resilience and vast tree mortality (Liu et al., 2019) and one study site that was used as evidence to measure the performance of the new methods. I also developed an R package that offers theses functionalities to explore early warning signals of forest mortality using spatio-temporal data.

## 2. Materials and methods

### 2.1. Overview

After describing the three case studies (Section 2.2), I explained the datasets, the steps used to only keep good quality pixels (Table 1, Section 2.3), and datasets acquisitions and preparations (Section 2.4). Although the previous studies with extensive field data reported the date of sudden tree mortality in the unhealthy ecosystems, I again used “breaks for additive seasonal and trend” (BFAST) to identify when the abrupt forest mortality occurred in each study site using MODIS satellite images (Section 2.5). I then created a subset of data for each study site between 2000 and two years prior to the abrupt change. Finally, I tested three methods, namely TEW (Section 2.8.1), SPT (Section 2.8.2), and the proposed approach by this study, i.e., STM and STG methods (Section 2.8.3), to test whether early warning signals of sudden forest mortality can be generated two years in advance.

**Table 1.** List of data products used in this study.

| Name | Unit | Product | Spatial resolution and sensor | Year |
| --- | --- | --- | --- | --- |
| Normalized difference<br>vegetation index<br>(NDVI) | - | MOD13A1 | 500-m<br>(MODIS) | 2000-2019 |
| Tree cover | % | MOD44B | 250-m<br>(MODIS) | 2001-2019 |
| Land cover | - | MCD12Q1 | 500-m<br>(MODIS) | 2001 |
| Palmer drought<br>severity index | - | TerraClimate | 0.5 degrees<br>(multiple satellites) | 2000-2019 |
| Digital elevation<br>model | meter | - | 90-m<br>(SRTM*) | - |

### 2.2. Study sites

I considered two forest ecosystems (i.e., Sequoia, and Yosemite) as the unhealthy case studies because they have been faced with severe tree mortalities, based on the reports of U.S. forest service (www.fs.usda.gov/detail/r5/forest-grasslandhealth/?cid=fsbdev3_046696, accessed date: 25.04.2020) and also Liu et al., (2019) (Fig. A1). I also selected Wood buffalo national park (hereafter Wood buffalo) as an evidence case in this study (Fig. A1). To the best of my knowledge, no sudden forest mortality has been reported for this national park. All three case studies are located in the national parks, where the exploitation of natural resources is prohibited. The spatial boundary datasets of Yosemite and Sequoia were obtained from the U.S. Government’s open data website (www.catalog.data.gov/dataset/national-parks, accessed date: 25.04. 2020), and the one for Wood buffalo national park was obtained from the Canada center for cadastral management’s cadastral datasets website (www.arcgis.com/home/item.html?id=2967061b2f48448bbd469906d971e401, accessed date: 25.04. 2020). To ensure the condition of the vegetation cover in the evidence study site has not changed beyond what is expected from natural processes such as phenology, I used the habitat intactness layer (www.intactforests.org, accessed date: 29.08. 2020) to identify the intact forest areas in Wood buffalo study site.

In all the case studies, the dominant forest type was evergreen needleleaf forest (Table 2). For the study time period of 2000–2019, the mean values of tree cover in Yosemite, Sequoia, and Wood buffalo were 47%, 51%, and 50%, respectively, and the mean values of the normalized difference vegetation index (NDVI) were 0.61, 0.68, and 0.66, respectively.

**Table 2.** Summary information for the study sites.

| Site | Dominant forests | Disturbance | Coordinate*<br>(latitude/longitude) | Number of MODIS pixels | Mean tree cover | Mean NDVI** |
| --- | --- | --- | --- | --- | --- | --- |
| Yosemite | Evergreen needleleaf | Drought and fungi attack*** | 37°75'74"N<br>119°63'28"W | 4052 | 47% | 0.61 |
| Sequoia | Evergreen needleleaf | Drought and fungi attack*** | 36°49'22"N<br>118°70'47"W | 1990 | 51% | 0.68 |
| Wood buffalo | Evergreen needleleaf | Not major disturbance is detected | 59°61'82"N<br>111°82'7"W | 10977 | 50% | 0.66 |

\* The coordinates correspond to the approximate center point of each study site.
\*\*Normalized difference vegetation index
\*\*\* *Phytophthora ramorum* fungi (Liu et al., 2019)

### 2.3. Datasets

#### 2.3.1. Slope

I used the digital elevation model (DEM) dataset, obtained from the shuttle radar topography mission (SRTM) with a 90-m spatial resolution, to calculate topographic slope values (Jarvis et al., 2008). I generated the slope values using the R “raster” package (the “terrain” function). I then aggregated the slope values spatially from 90-m to 500-m using the arithmetic mean function.

#### 2.3.2. Land cover map

To detect the type of forest within each study site, I used Land Cover Type 1 of the MODIS product (MCD12Q1, version 006), derived from the classification of the annual international geosphere-biosphere program (IGBP) (Channan et al., 2014) with a 500-m spatial resolution for the year 2001 (Sulla-Menashe and Friedl, 2019). I only selected forest types with tree cover greater than 30%, including evergreen needleleaf, evergreen broadleaf, deciduous needleleaf, deciduous broadleaf, mixed forest, and woody savanna classes (Friedl et al., 2010).

#### 2.3.3. Palmer drought severity index (PDSI)

To calculate the severity of regional drought conditions in each study site, I used the readily available monthly palmer drought severity index (PDSI) dataset in the TerraClimate product at 0.5-degree spatial resolution (Abatzoglou et al., 2014). TerraClimate has been estimated from several climatological data sources including WorldClim (Fick and Hijmans, 2017), CRU Ts4.0 (Jones and Harris, 2008), and Japanese 55-year Reanalysis (JRA55) (Agency, 2013). I downloaded the monthly PDSI data from the google earth engine platform (Gorelick et al., 2017) between 2000 and 2019. PDSI ranges between 10 (dry) and +10 (wet) which is estimated from temperature and precipitation data (Abatzoglou et al., 2018).

#### 2.3.4. Normalized vegetation difference index (NDVI)

I obtained NDVI from the latest version of the MODIS product, MOD13A1 (i.e., 006), which is improved through numerous upgrades in the algorithms (Didan, 2015). I downloaded NDVI data from the google earth engine platform (Gorelick et al., 2017) at a 500-m spatial resolution and a 16-day temporal resolution between 2000 and 2019. This product is computed from the atmospherically corrected bi-directional surface reflectance of the Red and NIR bands (0.620 to 0.670 μm, and 0.841 to 0.876 μm, respectively), ranging between −1 (bareland) and +1 (dense forest). The MOD13A1 data, by default, was masked for water, clouds, heavy aerosols, and cloud shadows. I used the standard quality tags information in the product to keep only the good quality and snow-free pixels. Since this product is not yet corrected for topographic-related issues, I used the topographic slope to exclude the pixels (from all the datasets used in this study) with the slope values greater than 25° (Fig. A2).

In this study, I used a vegetation-based indicator (i.e., NDVI), based on its successful performance as an ecosystem state variable to explore abrupt changes in the evergreen needleleaf forests (Liu et al., 2019; Rogers et al., 2018). Noteworthy, it has been shown that water-based indicators (e.g., normalized difference water index; NDWI) (Gao, 1996) provides more robust signals of ecosystem change, compared with vegetation-based indicators (e.g., NDVI) in the monitoring of forest ecosystems (Delbart et al., 2005; Wu et al., 2014). Nevertheless, since drought and water stresses, along with fungi attacks contributed to forest mortality (Liu et al., 2019), water-based indicators cannot truly represent the effects of such disturbances on vegetation cover (Fig. A3, Vescovi, 2017). Therefore, I selected NDVI to explore the state of study sites.

#### 2.3.1. Tree cover

I utilized NDVI and tree cover together to get an overview of both the understory and overstory changes in the case studies. The NDVI product may correspond to the total NDVI of a pixel, representing both the understory and overstory vegetation, while tree cover represents overstory in a forest. I used vegetation continuous fields (VCF), version 006, of MODIS to obtain the tree cover dataset that ranges between 0 (no tree cover) and 100 (dense tree cover). I downloaded the yearly product of tree cover from the google earth engine platform (Gorelick et al., 2017) at a 250-m spatial resolution between 2001 and 2019. This product was prepared using monthly composites of Terra MODIS 250- and 500-meters land surface reflectance data, including all the bands (i.e., one to seven), and land surface temperature. I kept only the good quality tree cover pixels (i.e., estimated at clear sky) using the quality information available in this product.

### 2.4. Dataset preparations

To ensure having enough high-quality NDVI observations in the study sites, first, I aggregated the data temporally from a 16-day to a monthly resolution using the mean function. Given the importance of properly considering the distribution of missing values over time, I obtained the temporal pattern of missing values in each study site (Fig. A4). The results showed that the number of available pixels in the winter-time of 2014 was significantly higher (due to the low precipitation) than those of the other years in Yosemite and Sequoia. Having more observations in the winter-time in 2014 can falsely show abrupt decreases in the NDVI time series and may increase the risk of false alarming in identifying early warning signals. Although interpolation methods (e.g., linear model and moving average) may be used to fill the missing values, it may lead to a high risk of false alarms as it increases the autocorrelation values (Dakos et al., 2012a). Therefore, I excluded the data between December and March (four months) from the time series of NDVI in all the case studies. This four-month time window was selected based on data quality information of NDVI (Fig. A4).

### 2.5. Breaks for additive seasonal and trend method (BFAST)

I identified the date of the largest significant abrupt change in the time series of NDVI for each study site using the “breaks for additive seasonal and trend” (BFAST) method (Verbesselt et al., 2012). A breakpoint, here, refers to the largest significant change in the time series of NDVI between 2000 and 2019 that could represent forest mortality.

To examine the breakpoint occurrence, first, I extracted global temporal NDVI values (i.e., mean NDVI values of all the pixels in each month) by aggregating the data using a mean function. Then, I decomposed the NDVI time series into the trend and seasonal components (Verbesselt et al., 2010). The breakpoints, then, were calculated using the BFAST method (Verbesselt et al., 2010) which iteratively fits a piecewise linear trend and a seasonal model to the time series data (from time points 1 to n) using:

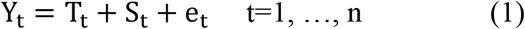

where Y_t_, is the observed data at time point t, T_t_, is the trend component, S_t_, is the seasonal component, and e_t_, is the remainder component (Verbesselt et al., 2010). It is assumed that T_t_ is a piecewise linear function with m+1 segments. Each segment has a specific slope and intercept. Thus, there are m breakpoints 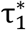,…, 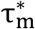, such that:

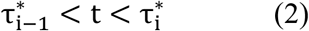

where i = 1,…, m.

In this study, I used breakpoints of a trend at 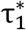,…, 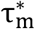 that can be estimated using the residual-based moving sum test, and were assessed by minimizing a Bayesian information criterion (BIC) from the seasonally adjusted data Y_t_ – Ŝ_t_, where Ŝ_t_ is found by the seasonal and Trend decomposition using the loess (STL) method (Cleveland et al., 1990). STL is a filtering procedure for decomposing a seasonal time series into trend, seasonal, and residual components using a sequence of applications of the loess smoother.

The trend between breakpoints is given as:

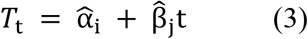

where and *i* = 1,…, m and intercept 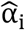 and slope 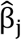 can be used to calculate the magnitude of the abrupt change and slope of the gradual change between detected breaks points (Venables and Ripley, 2002; Verbesselt et al., 2010).

In this analysis, the number of breakpoints per time series was restricted to one, which was the largest significant breakpoint. After finding the breakpoint in the time series, I only considered the time series of each study site temporally between the year 2000 and two years before the timing of abrupt change occurrence. The two-year time window was also used in a previous study (Liu et al., 2019). I also mapped the trend of changes in NDVI until the time of the breakpoint in each study site pixelwise (results in Fig. 4).

**Fig. 1.**
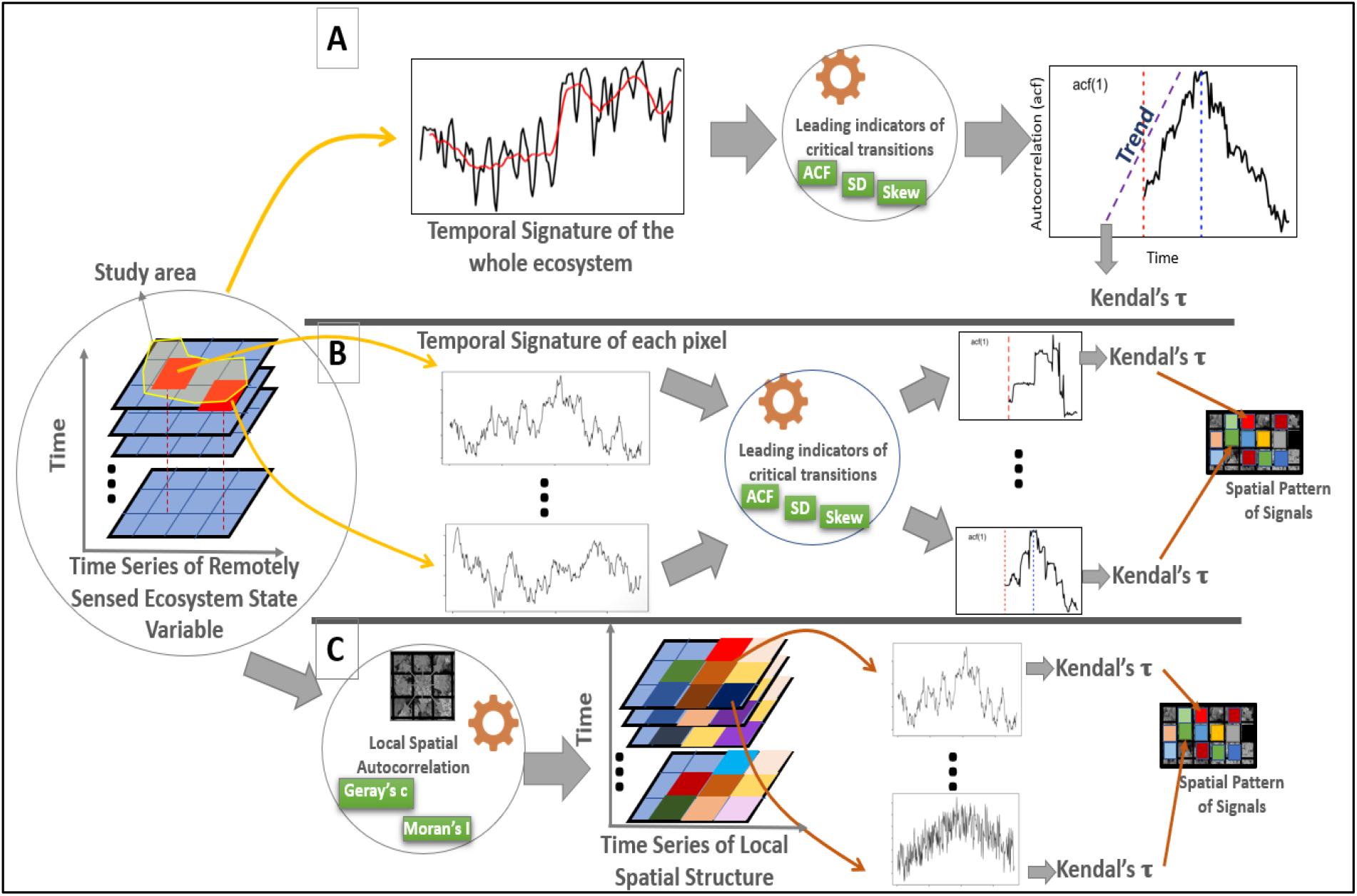
Conceptual diagram representing different approaches used in this study to measure early warning signals of forest mortality; A) analysis of time series represents the dynamics of the whole ecosystem state variable, where the temporal signature is averaged over the whole ecosystem and after that leading indicator of critical transitions has been measured; B) following the approach provided in (A) but applied to the time series obtained from each pixel of a spatio-temporal time series that generates a spatial pattern of early warning signals; C) a new approach proposed in this study that measures local spatial autocorrelation (using local Moran’s *I* and local Geary’s *c*) at each time and generates early warning signals from time series of local spatial autocorrelation values at each pixel.

**Fig. 2.**
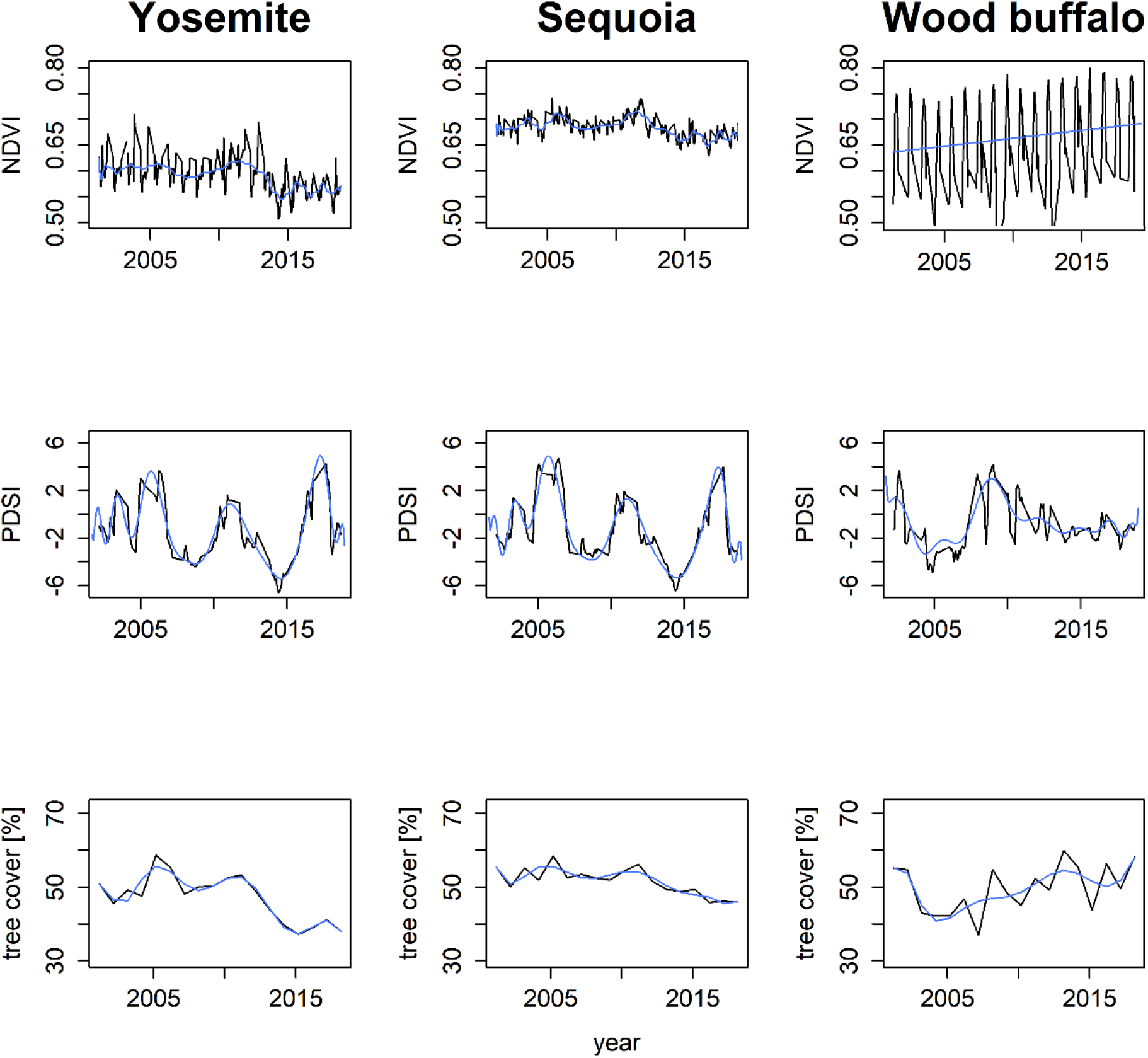
The time series of normalized difference vegetation index (NDVI), palmer drought severity index (PDSI), and tree cover in Yosemite, Sequoia, and Wood Buffalo national park (Wood buffalo) study sites between 2001 and 2019. In each graph, the blue line represents the smoothed time series fitted using a generalized additive model.

**Fig. 3.**
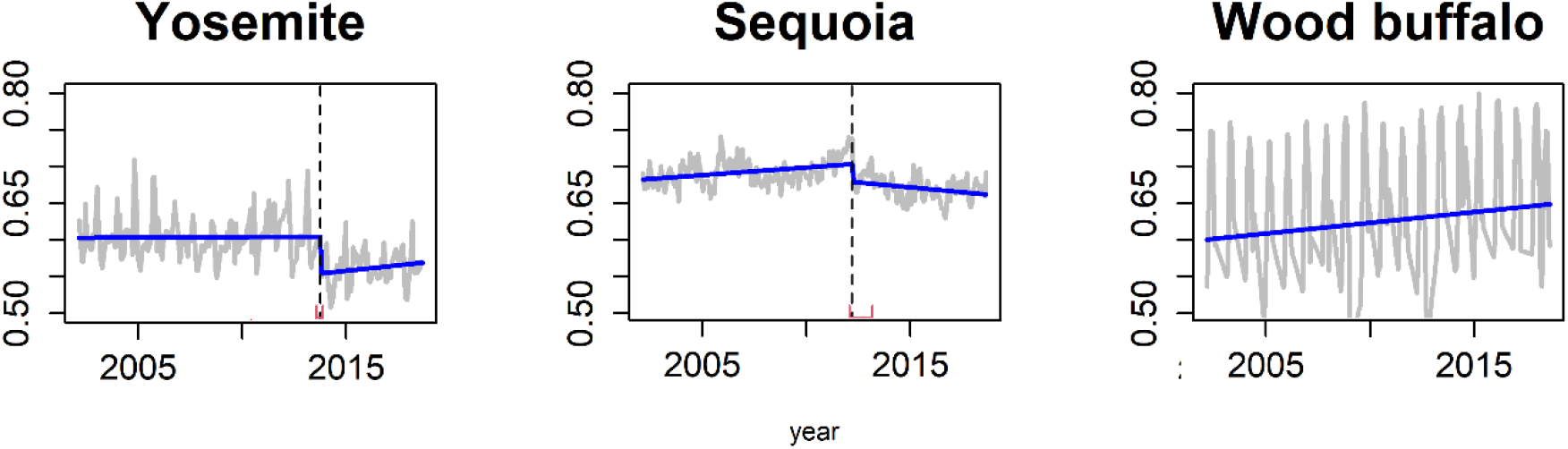
Breakpoints in time series of normalized difference vegetation index (NDVI) in the Yosemite, Sequoia, and Wood buffalo national park (Wood buffalo) case studies based on the breaks for additive seasonal and trend (BFAST) method. The gray lines represent actual data, the blue lines represent the fitted model. The dashed vertical lines indicate the date of the breakpoint occurrence, and the red lines represent the confidence interval of the timing of the breakpoints.

**Fig. 4.**
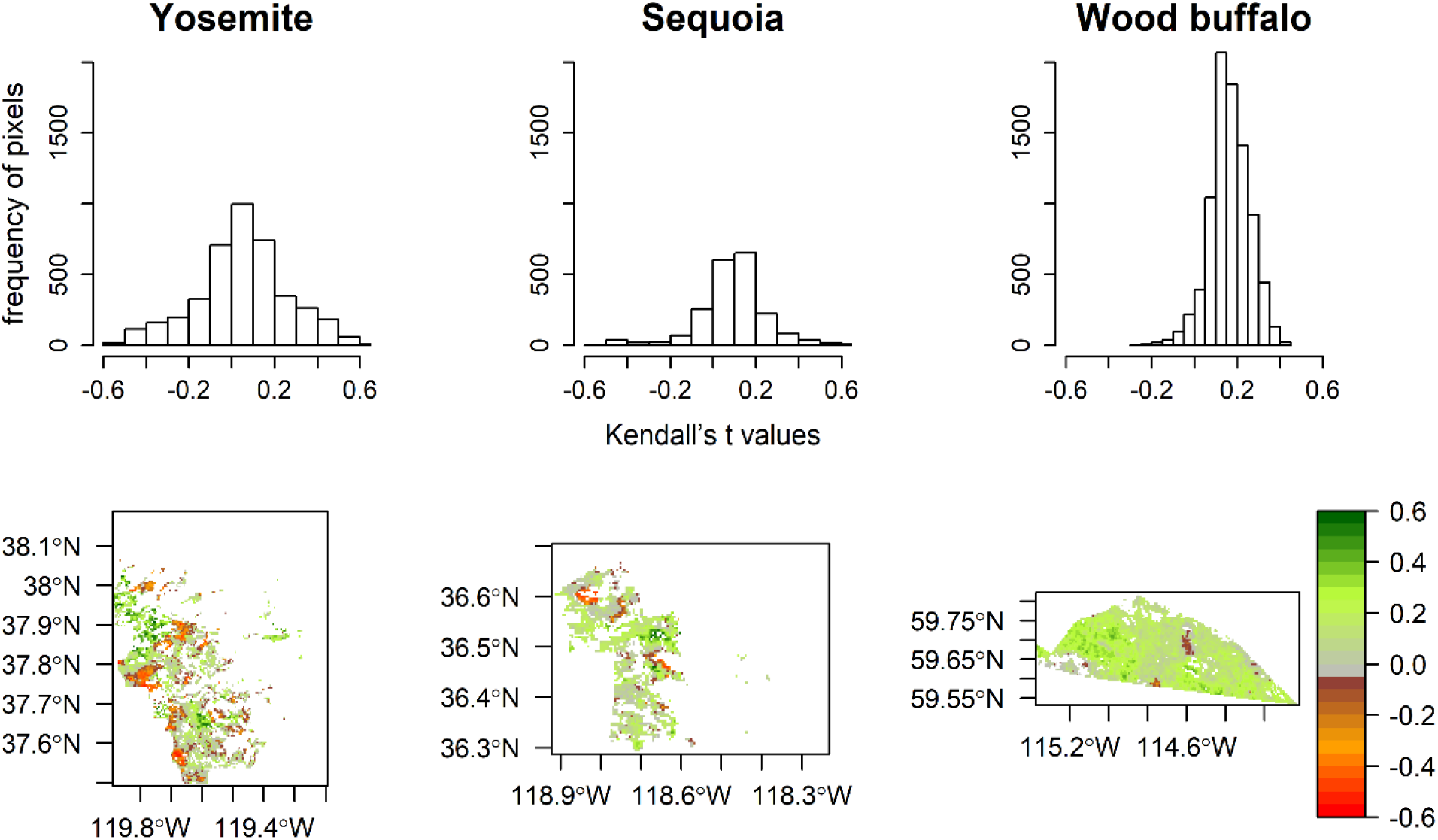
The magnitude of the trend in the NDVI time series at each spatial location (pixel) between 2000 and 2019, calculated using Kendall’s τ, in different case studies. The first row shows the histogram of the trend values over different pixels, and the second row represents the spatial distribution of the trend values.

### 2.6. Approaches to detect early warning signals

I tested three different approaches to identify early warning signals, including TEW, SPT, and the new proposed approach uses STM and STG. The TEW uses a temporal signature (time series) of the whole ecosystem to which the early warning signals are applied. SPT applies TEW on all the high-quality pixels within each study site and generates the spatial distribution of the early warning signals. STM and STG perform local spatial autocorrelation statistics pixelwise in time and then estimates the strength of early warning signals has been observed (Fig. 1).

#### 2.6.1. Temporal early warning signals (TEW) and Spatial pattern of temporal early warning signals (SPT)

Statistical indicators, such as increased autocorrelation in the temporal signature of an ecosystem state variable (in this study, NDVI), are inferred as warning signals of reduced resilience (Scheffer et al., 2012). The lag-1 autocorrelation is a commonly used indicator for this purpose calculated by:

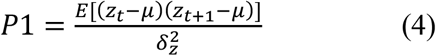

where *E* is expected value, *μ* is mean, and *δ* the variance of a variable *z_t_* (Bartholomew et al., 1971).

I first extracted NDVI time series averaged over the whole ecosystem. Next, I detrended the time series to eliminate the effects of non-stationary using the first differencing method, where the Augmented Dickey-Fuller test (Banerjee et al., 1993) confirmed the non-stationary condition is eliminated from the data using the first differencing method. To calculate the lag-1 temporal autocorrelation I used a rolling window of 50% (to measure autocorrelation using a subset of data located within the rolling window), which is not very short or very long. Then, I calculated the trend of changes in the autocorrelation of the NDVI time series using the non-parametric Mann Kendall test (named as Kendall’ τ, hereafter). A Kendal’s τ of zero implies no significant trend of changes, and the value of +1.0 or −1.0 implies a perfect positive or perfect negative (inverse) trend, respectively.

The spatial pattern of temporal early warning signals, then, was calculated by applying TEW pixel-wise and then assigning the results of Kendal’s τ test to each pixel. In other words, I applied TEW on all the high-quality pixels within the geographical extents of each study site to generate the spatial distribution of the corresponding signals.

#### 2.6.2. Temporal analysis of local spatial structure (STM and STG)

To calculate the STM and STG, first, I used two local spatial autocorrelation statistics (i.e., local Moran’s *I* and local Geary’s *c*) and applied them to the image of NDVI at each time step.

Second, I extracted the values of the local spatial autocorrelation at each pixel between 2000 and two years prior. Third, I smoothed the extracted values using a moving average, with the size of three, to reduce the effects of, for example, precipitation occurrence that can contribute to autocorrelation values (Verbesselt et al., 2016). Finally, I calculated the trend of changes in the time series of the local spatial autocorrelation at each pixel using the Kendall’ τ test. I used Kendall’s τ to calculate the monotonic trend in the time series of local Moran’s *I* or local Geary’s *c*. For the trend analysis, sometimes, the deseasonalisation step is taken to avoid failing the stationary assumption in the data. However, this procedure is not followed in this study, since in the context of environmental time series, it is expected that the real trend of changes cannot greatly be affected by seasonality (Hipel and McLeod, 1994).

I estimated the local Moran’s *I* at site *i* using (Anselin, 1995):

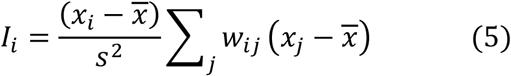

where *I_i_* is local Moran’s *I* for site *i*, *w_ij_* is a spatial weight for an observation *j* (often binary, i.e. 1 for neighboring locations and 0 elsewhere), 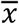 and *s*^2^ are the global sample mean and variance of the observations, respectively, and *x_i_*, and *x_j_* are the observed values at site *i* and site *j* (i.e. the neighborhood of site *i*), respectively. Positive values of *I_i_* indicate a cluster of similar values around site *i*, and negative values indicate that the neighbor values are dissimilar to site *i*.

Local Geary’s *c* statistic at site *i* is (Anselin, 1995):

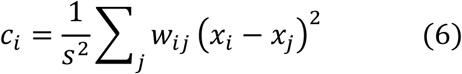

with the same variables as local Moran’s *I*. This statistic quantifies a standardized squared distance between the values at site *i* and the neighbor locations *j*. High values of *c_i_* indicate substantial differences between site *i* and its neighbors, and low values of *c_i_* indicate that site *i* is similar to its neighbors. Local Moran’s *I* measures deviations of locations from the global mean. Local Geary’s *c* summarizes differences between a site and the values of its neighboring sites.

All statistical analyses and visualizations were performed in R statistical software (Somanathan et al., 2004), and Google Earth Engine (Gorelick et al., 2017). I used different R packages, including “raster” (for basic raster analysis) (Hijmans, 2015), “rts” (for time series analyses) (Naimi, 2016), and “bfast” (for the BFAST analysis) (Verbesselt et al., 2012).

## 3. Results

### 3.1. Overall ecosystem changes

The PDSI time series showed that Yosemite and Sequoia were faced with two major regional drought conditions spanning 2007–2009 and 2014–2016 that can be reflected in NDVI. In Wood buffalo study site, NDVI showed regular seasonality and PDSI revealed severe drought conditions in 2005. For the entire Yosemite and Sequoia, the fraction of the tree cover was decreased after 2012, that was not recovered afterward. The timing of the irreversible tree cover and NDVI declines was consistent with the severe drought condition shown by the PDSI values in Yosemite and Sequoia. The tree cover in the Wood buffalo study site showed some fluctuations, recovered after each tree cover loss.

Yosemite and Sequoia showed their largest significant breakpoints in NDVI in 2013 and 2011, respectively, while Wood buffalo showed no breakpoint (Fig. 3). The changes of NDVI after the breakpoint occurrence demonstrated a slight decrease in the mean NDVI in the Yosemite and Sequoia case studies.

### 3.2. NDVI changes

The results showed a significant decrease in the trend of NDVI in some forest patches in Yosemite and Sequoia, whereas no major change was observed in the Wood buffalo study site (Fig. 4). The histogram of the NDVI trend (Kendall’s τ) revealed that the range of the trend values was between −0.6 and 0.6 in Yosemite and Sequoia, while it was between −0.3 and 0.4.5 in Wood buffalo (Fig. 4).

### 3.3. Temporal analysis of early warning signals (TEW)

In the Yosemite study site, autocorrelation showed a significant decrease (Kendall’s τ = −0.26) (Fig. 5) In the Sequoia study site, the general autocorrelation mirrored a significant decreasing trend (Fig. 5, Kendall’s τ = −0.52). In the Wood buffalo study site, the trend of autocorrelation changes showed a higher magnitude (Fig. 5, Kendall’s τ = −0.15) compared with the other case studies, where autocorrelation was more erratic than the other case studies.

**Fig. 5.**
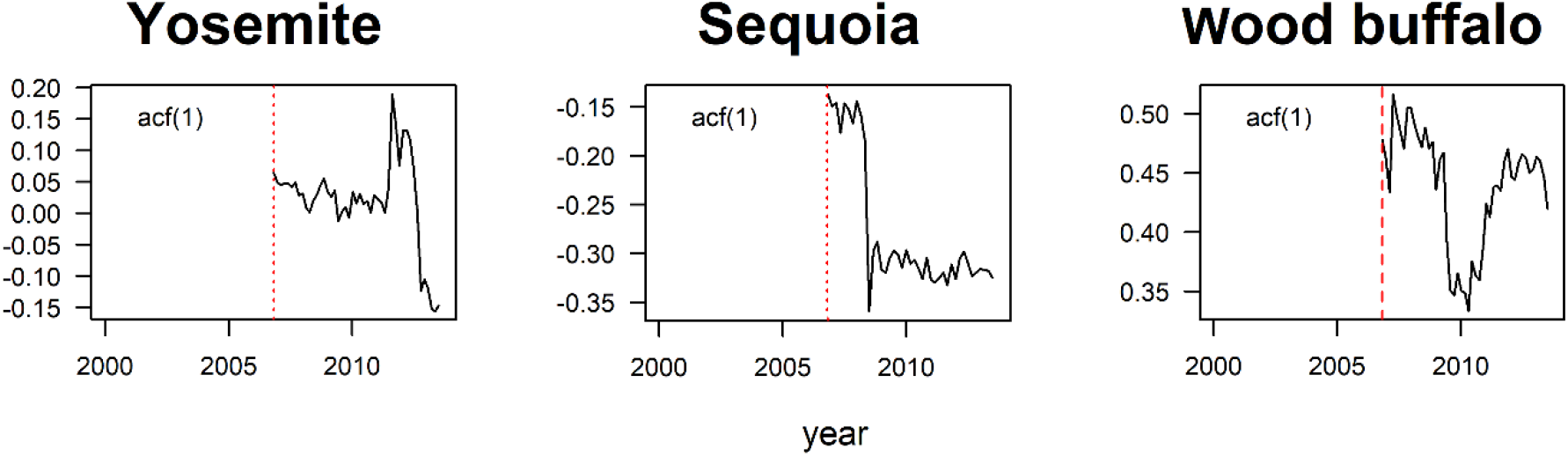
Lag-1 autocorrelation, acf(1), of the NDVI time series. In each graph, the y-axis represents the autocorrelation values, and the x-axis represents time (year), ranging between 2000 and two years before the abrupt change identified in each study site. The red dotted lines mark the size of the rolling window used to calculate acf(1).

### 3.4. The SPT, STM, and STG results

The magnitude of changes in the local spatial autocorrelation, obtained from the trend analysis of the values at each pixel until two years prior to the identified abrupt change, varied considerably pixel-wise in Yosemite and Sequoia (Fig. 6). In some patches of forests, where NDVI was declined (Fig. 4), the results highlighted increases in STG values and decreases in STM values (Fig. 6). SPT values varying over space in study sites. The shape of histograms suggests that the values of STM and STG were normally distributed, while the distribution of values in Wood buffalo is more uniform or biased toward the negative or positive values (Fig. 7).

**Fig. 6.**
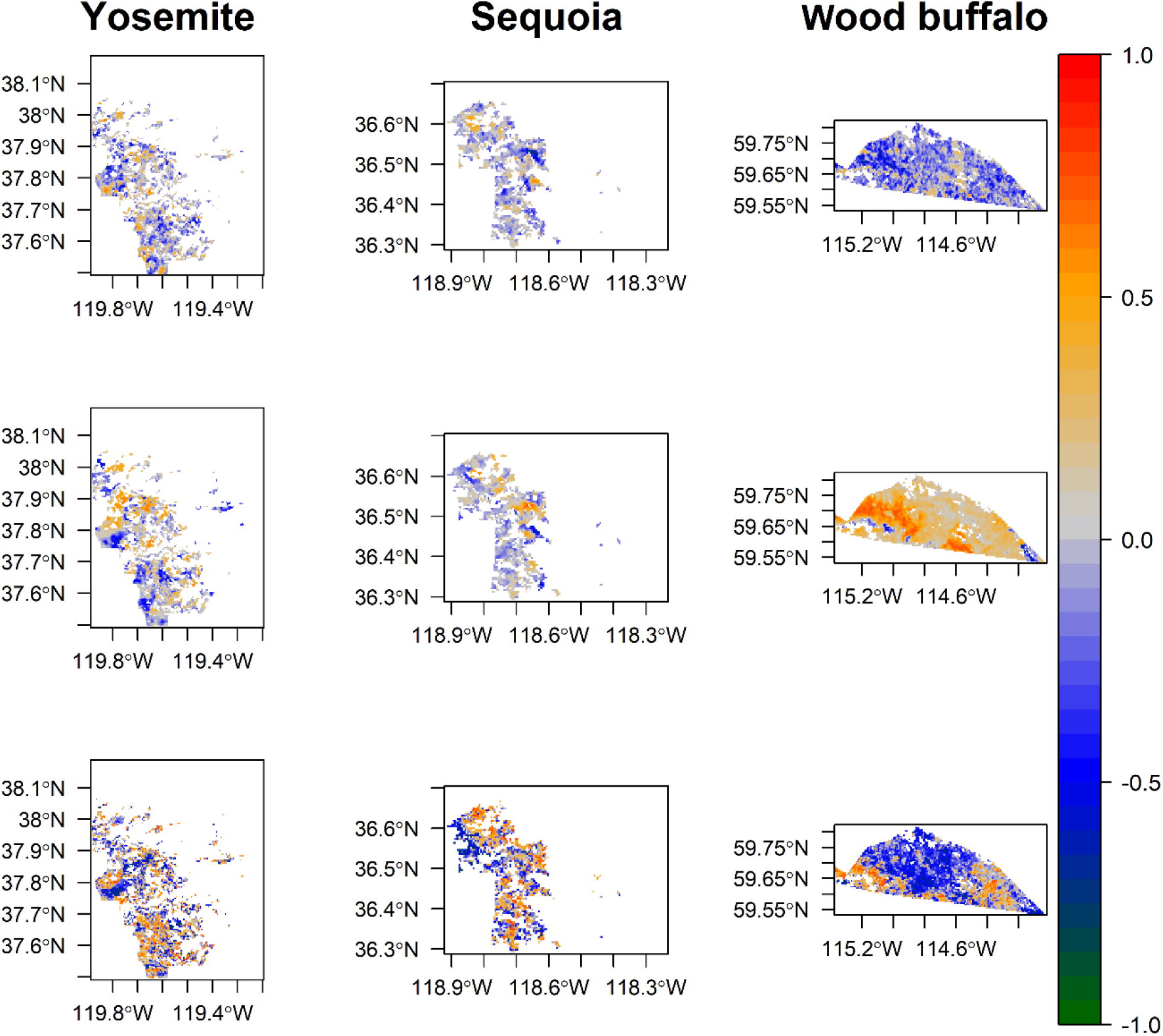
Early warning signals of forest mortality in the Yosemite, Sequoia, and Wood buffalo national park (Wood buffalo) case studies. The pixel values represent the magnitude of trends (Kendall’s τ); the first row is based on the time series of local Geary’s *c*, the second row is based on the time series of local Moran’s *I*, and the third row is based on the temporal analysis of early warning signals (lag-1 autocorrelation) at each pixel. In each map, the x-axis represents longitude, and the y-axis represents latitudes.

**Fig. 7.**
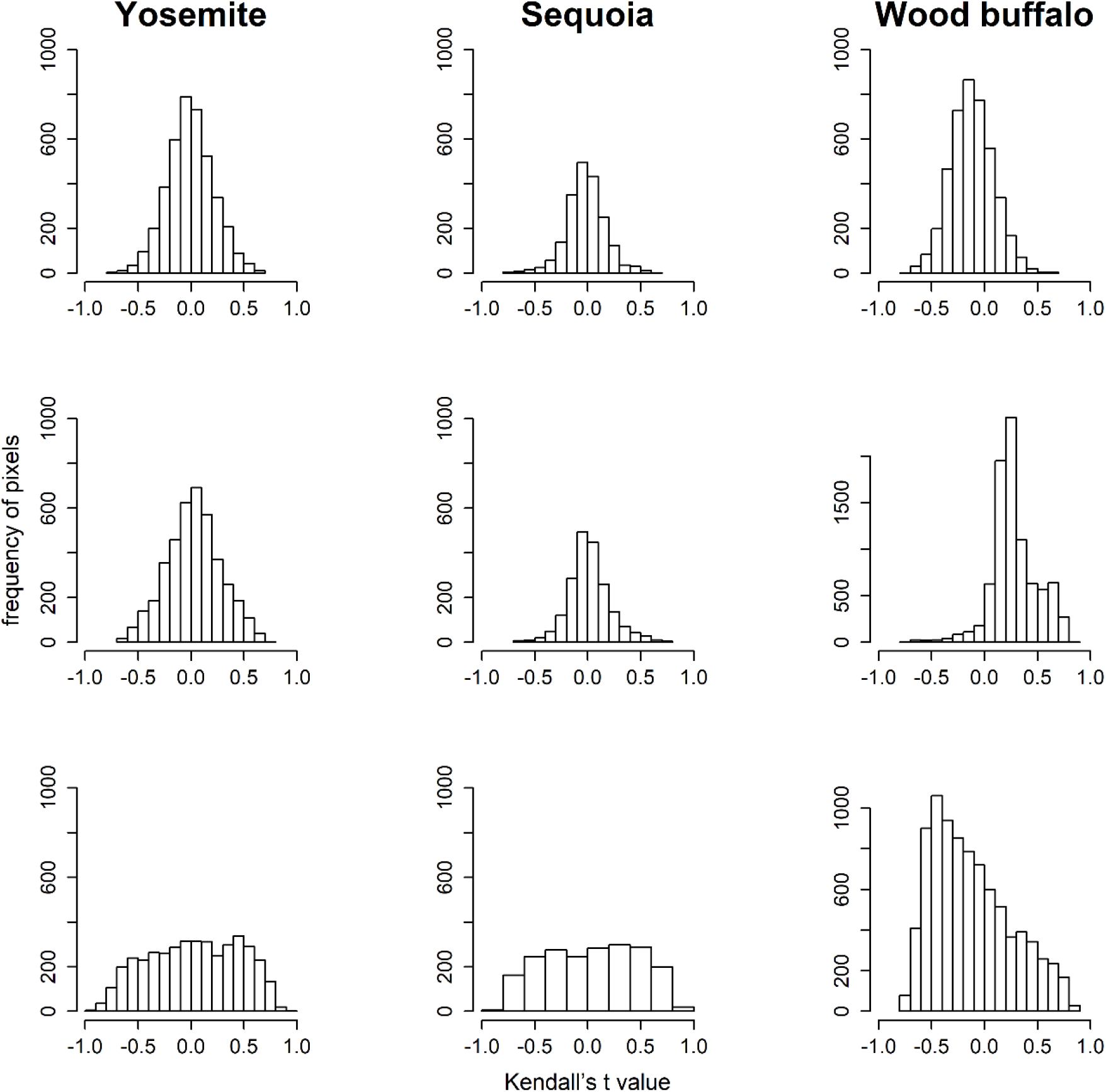
Histogram of Kendall’s τ values over different pixels (Fig. 6) in the Yosemite, Sequoia, and Wood buffalo national park (Wood buffalo) case studies. The first row is related to the local Geary’s *c*, the second row is related to the local Moran’s *I*, and the third row is related to lag-1 autocorrelation (temporal analysis of early warning signals).

### 3.1. R package

So far, two main R packages have been developed to support the monitoring of ecosystem dynamics, including “earlywarnings” that assist TEW (Dakos et al., 2012a) and “spatialwarnings” that contribute to the furtherance of space for time analysis (Kéfi et al., 2014). In this study, I introduced a new R package, called “stew”, that facilitates to analyze spatio-temporal data and explore early warning signals of a critical transition. The R package was designed (1) to import time series of satellite images, (2) to measure local Geary’s *c* and local Moran’s *I* and generate time series of local spatial structure, and (3) to estimate the trend of autocorrelation changes using Kendal’s τ. Besides, the “stew” R package ameliorate (4) pixel wise statistical analysis using, for example lag-1 autocorrelation, standard deviation) to generate a spatial distribution of early warning signals. These analyses were benefitted from the possibility of using different sizes of a rolling window and different methods of detrending (i.e., first-differencing, loess, and gaussian). Moreover, the software (5) automatically calculate STM and STG methods and provides a set of functions to (6) visualize and export the output of the analysis as a spatial map.

## 4. Discussion

This study contributes to an understanding of identifying the early warning signals before climate-induced forest mortality occurrence using remotely sensed spatio-temporal indicators. The results of this study showed the successful performance of STM and STG in generating early warning signals of forest mortality (Fig. 6–7), which is consistent with the previous report about the adequacy of spatial indicators to identify critical transition (Dakos et al., 2015).

### 4.1. TEW and SPT approaches

The results of TEW between 2000 and the two years prior to forest mortality showed false negatives, i.e., no early warning signals while it should alarm, in Yosemite and Sequoia by Kendall’s τ of −0.26 and −0.52 respectively (Fig. 5). The signals in TEW analysis vanish since temporal analysis measures the state of an ecosystem in time through quantifying the variation of an ecosystem state variable averaged over the whole ecosystem. By aggregating the data, the negative and positive values of signals may cancel each other out, resulting in false-negative alarms. Liu et al (2019) also illustrated false-negative results in forests located in California, where Yosemite and Sequoia are located. According to this study, one reason behind the false-negative results is the distribution of different forest types with varying resilience across the study area. For example, some trees are more adaptive to drought conditions, which can experience an abrupt change later than other trees (Liu et al., 2019). Although TEW failed to provide an accurate alarm for unhealthy study sites, it provided a true negative alarm in the Wood buffalo study site by Kendall’s τ of −0.15 (Fig. 2–4).

The SPT analysis showed varying results with a high rate of the false-negative alarms (Fig. 6) when one visually tracks forest mortality locations (Fig. 4). Especially for the evidence study site (Wood buffalo), it showed false-positive alarm i.e., an early warning signal while it should not alarm, with Kendall’s τ of more than 0.5 in majority of pixels (Fig. 7). Such false alarms may occur due to the sensitivity of TEW analysis to, for example, pre-processing, or differences in vegetation composition in a forest. Liu et al (2015) argued that the false positive might be related to some forest pixels that are unhealthy without tree mortality phenomenon.

### 4.3. STM and STG approaches

Increases in STM and decreases in STG (Fig. 6) where NDVI decreased (Fig. 4), refers to local spatial autocorrelation reductions. Declines in local spatial autocorrelation happen when a high level of ecosystem fragmentations is occurred (Kowe et al., 2019). As forest fragmentation continues, meaning that forests increasingly are divided into separated portions with smaller and more isolated patches, the critical transition is expected to happen (Taubert et al., 2018). Given the observed local spatial autocorrelation reductions using STM and STG, both methods provided true positive alarms in unhealthy study sites. The opposite scenario is true for the evidence ecosystem, where the STM and STG provided true negative alarms.

From a methodological point of view, one reason behind the acceptable performance of STM and STG compared with temporal analysis is that the choice of the window size, which may significantly alter the results of TEW approaches (Emmanuel and Tamga, 2019), are not applicable for STM and STG. Furthermore, TEW should be followed by trend and seasonality decompositions from the data prior to the analysis, however, there is no golden rule on the pre-processing steps, which may invite some uncertainties into the results. For instance, Verbesselt et al., (2016) argued that the results of TEW may be sensitive to the detrending methods. As a solution, Verbesselt et al., (2016) used an ensemble of six different detrending methods, though especially in evergreen forests, the imperfect seasonal detrending of foliage variation could affect the results of the TEW (Verbesselt et al., 2016). Nevertheless, STM and STG methods may be more robust to identify early warning signals of forest mortality since they do not require the use of a rolling window, which makes the mothed less sensitive to pre-processing.

One advantage of STM and STG compared with temporal analysis is that they are less expensive for larger-scale applications, e.g., real-time global forest monitoring. Moreover, the proposed approaches, opposite to spatial analysis (e.g., methods proposed by Kéfi et al., 2014), are less sensitive to the type of input data (categorical or continuous) that can decrease the uncertainties related to the classification of satellite images with a coarser resolution (i.e., MODIS) and quantifying the signals of impending critical transition.

### 4.4. Uncertainties and limitations

I tested the performance of TEW, SPT, STM, and STG approaches in only the evergreen needleleaf forests in three limited study sites. For a broader application, the performance of the methods on different locations with different forest types and climate conditions should be investigated. In addition, due to the data medium spatial resolution, differences among the species were not studied in detail, which could be a hindrance against the full achievement of the chief objectives of this article. Furthermore, I performed the analysis in national parks, where the direct effects of human influences are supposed to be limited. It suggested that the method should be further examined and validated in other ecosystems.

It has been shown that the success of detecting early warning signals is crucially dependent on the spatial resolution of satellite images (Nijp et al., 2019). In the coarse resolution images, the signals can be compensated by having different overstory and forest floor optical properties (i.e., one pixel represents the optical properties of different species and soil chokecherries). On the other hand, using satellite images with a fine spatial resolution may increase the computational efforts and decrease the applicability of the method in monitoring forests on a large scale, e.g., a global scale. In this study, I showed the successful application of a MODIS product with a not too coarse or fine spatial resolution. The high-temporal resolution of MODIS products can facilitate the application of such methods for global studies (Fig. 7). Nevertheless, optical satellite images are subjected to uncertainties due to cloud impacts on the acquired data. Although we carefully preprocessed the data to produce reliable results, the usage of a new generation of radar mission sentinel data (sentinel 1 and 3), which provides cloud-free datasets, is recommended for future studies.

## 5. Conclusion

I used a spatio-temporal analysis of satellite images to test the performance of different methods, detecting early warning signals in two unhealthy forest sites and one evidence forest site. I showed the ScTM and STG approaches outperformed TEW and SPT to provide early warning signals in two years prior to an abrupt change in the study sites. This confirms that spatial indicators may be more reliable than a temporal analysis, which is consistent with the previous reports. I also developed an R package that supports analysis of ecosystem dynamics, called “stew”.

## Acknowledgments

I would like to thank Dr. Miina Rautiainen and Dr. Babak Naimi for their insights and comments on this article.

## Supplementary information

**Fig. A1.**
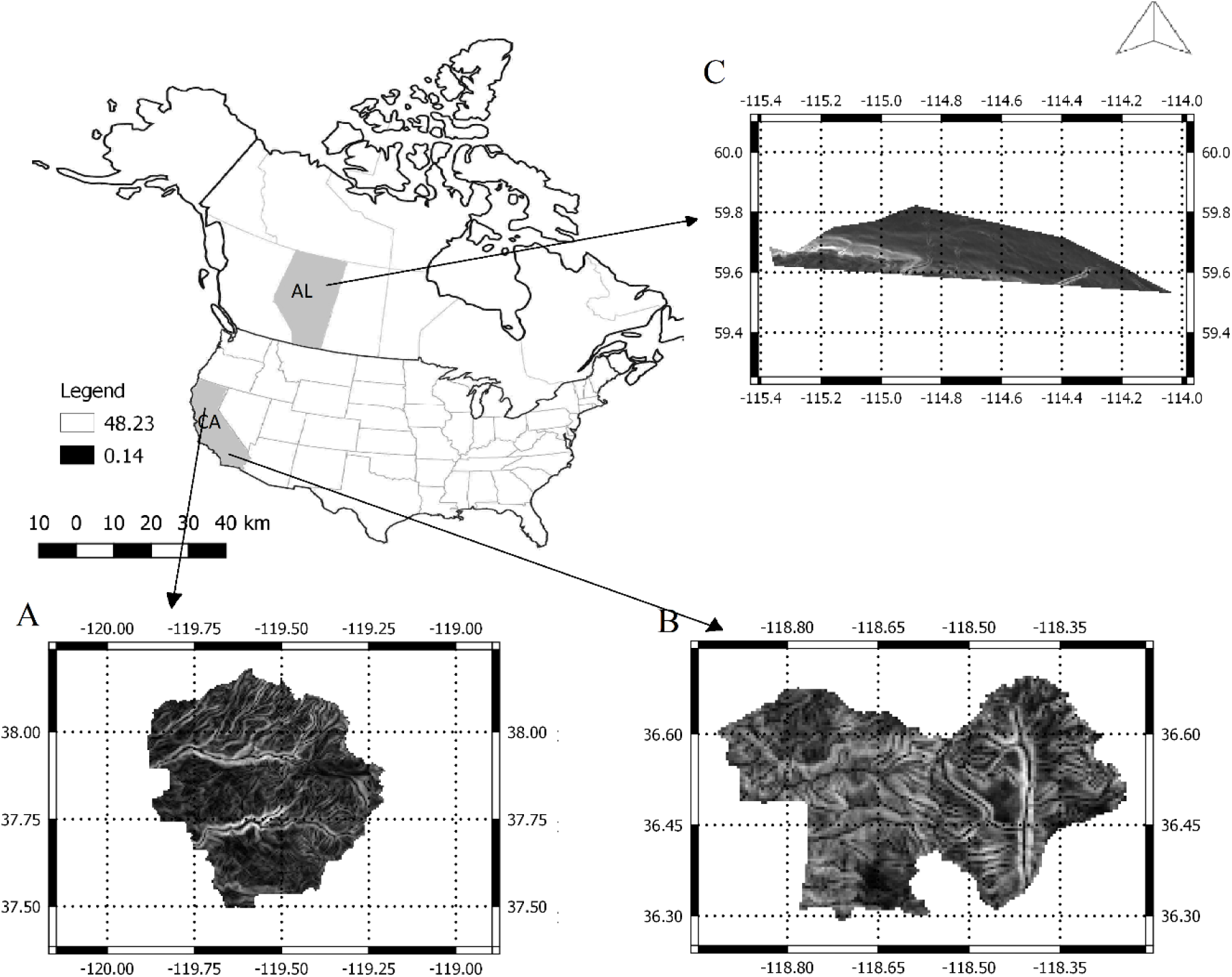
The location of each study site. A: Yosemite study site in California (Ca) in America, B: Sequoia, C: Wood buffalo national park in Alberta (Al), Canada.

**Fig. A2.**
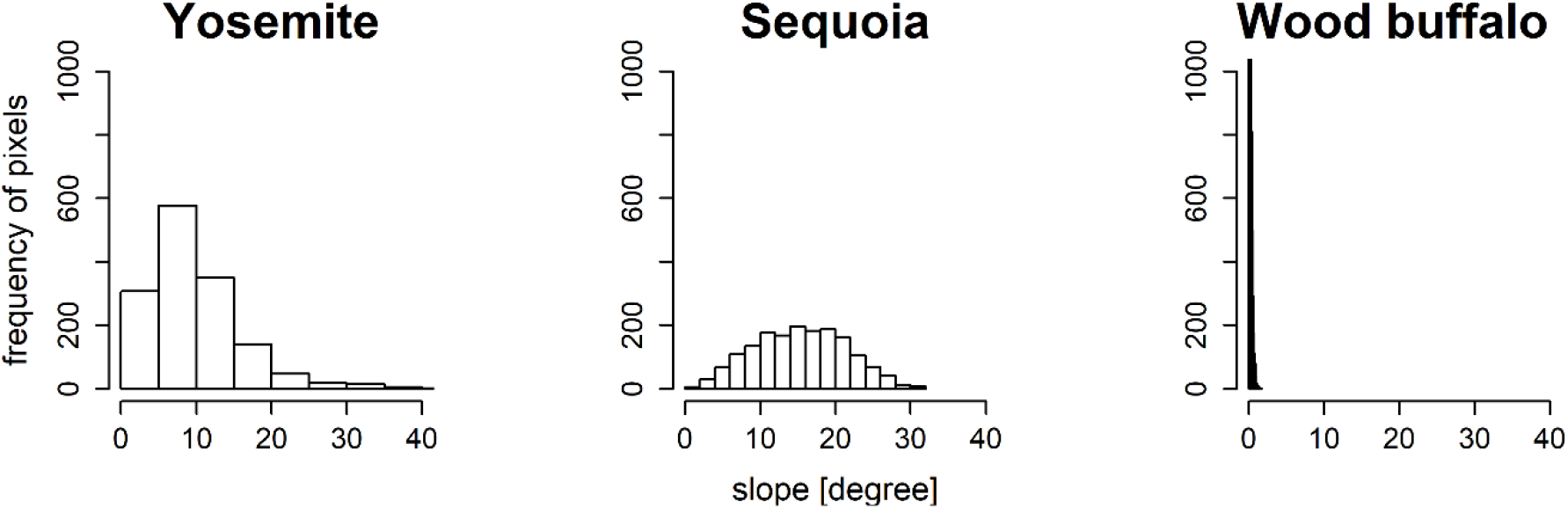
The frequency of forest pixel counts within each study site, including Yosemite, Sequoia, and Wood buffalo national park (Wood buffalo). In each graph, the y-axis represents the frequency of pixels and the x-axis represents slope.

**Fig. A3.**
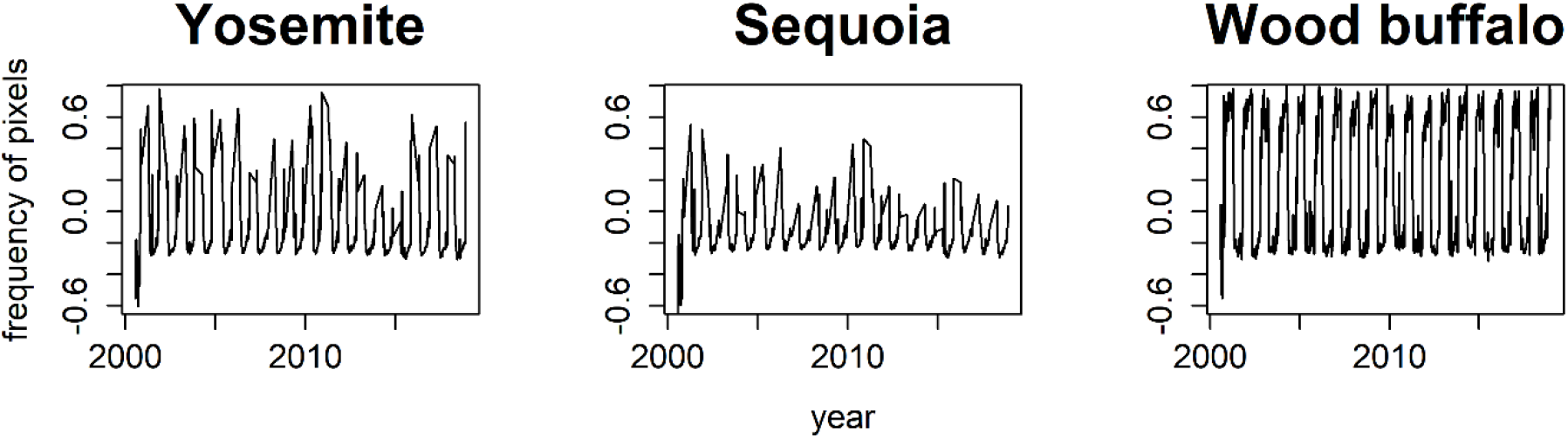
Time series of normalized difference water index (NDWI) between 2000 and 2019 in Yosemite, Sequoia, and Wood buffalo national park (Wood buffalo). In each graph, the y-axis represents the NDWI and the x-axis represents time.

**Fig. A4.**
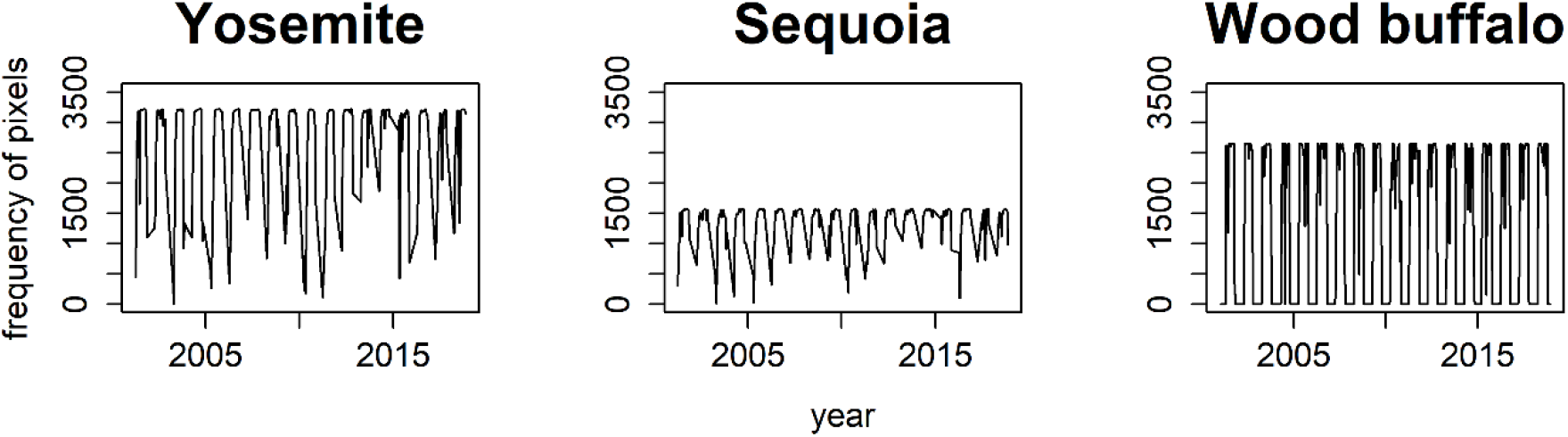
Percentage of high-quality normalized difference vegetation index (NDVI) retrievals in the Yosemite, Sequoia, and Wood buffalo national park (Wood buffalo) study site between 2000 and 2019 based on the NDVI quality data set (MCD12Q1).

